# Fluorinated oil-surfactant mixtures with the density of water: artificial cells for synthetic biology

**DOI:** 10.1101/2021.05.17.444437

**Authors:** Roberto Laos, Steven Benner

## Abstract

There is a rising interest in biotechnology for the compartmentalization of biochemical reactions in water droplets. Several applications, such as the widely used digital PCR, seek to encapsulate a single molecule in a droplet to be amplified. Directed evolution, another technology with growing popularity, seeks to replicate what happens in nature by encapsulating a single gene and the protein encoded by this gene, linking genotype with phenotype. Compartmentalizing reactions in droplets also allows the experimentalist to run millions of different reactions in parallel. Compartmentalization requires a fluid that is immiscible with water and a surfactant to stabilize the droplets. While there are fluids and surfactants on the market that have been used to accomplish encapsulation, there are reported concerns with these. Span® 80, for example, a commonly used surfactant, has contaminants that interfere with various biochemical reactions. Similarly, synthetic fluids distributed by the cosmetic industry allow some researchers to produce experimental results that can be published, but then other researchers fail to reproduce some of these protocols due to the unreliable nature of these products, which are not manufactured with the intent of being used in biotechnology. The most reliable fluids, immiscible with water and suitable for biochemical reactions, are fluorinated fluids. Fluorinated compounds have the peculiar characteristic of being immiscible with water while at the same time not mixing with hydrophobic molecules. This peculiar characteristic has made fluorinated fluids attractive because it seems to be the basis of their being biologically inert. However, commercially available fluorinated fluids have densities between 1.4 to 1.6 g/mL. The higher-than-water density of fluorinated oils complicates handling of the droplets since these would float on the fluid since the water droplets would be less dense. This can cause aggregation and coalescence of the droplets. Here, we report the synthesis, characterization, and use of fluorinated polysiloxane oils that have densities similar to the one of water at room temperature, and when mixed with non-ionic fluorinated surfactants, can produce droplets encapsulating biochemical reactions. We show how droplets in these emulsions can host many biological processes, including PCR, DNA origami, rolling circle amplification (RCA), and Taqman^®^ assays. Some of these use unnatural DNA built from an Artificially Expanded Genetic Information System (AEGIS) with six nucleotide “letters”.

## Introduction

There is a growing interest in the biochemical sciences for the compartmentalization or miniaturization of reactions [1] [2]. When conducted in water-in-oil droplets, compartmentalization allows experimentalists to run millions of reactions in parallel, creating ample opportunity to analyze the vast diversity contained within a biochemical sample. One can amplify an isolated gene in a digital PCR or explore the vast protein sequence space in directed evolution experiments [3]. Approaches such as directed evolution and/or high throughput screening require sequence diversity. These kinds of experiments require to isolate variants of a biomolecule. In some cases, enzymes can be isolated with their encoding genes and can be, considered by some, as forming a sort of “artificial cell”. One way to achieve this compartmentalization and screening is with the use of water-in-oil droplets [4]. These “artificial cells” [5] [6] and other compartments have several applications, including drug delivery [7-9], enzyme directed evolution [10, 11] [12, 13] [1] [14], and models for the origin of life [15-19]. While water-hydrophobe-water systems are analogs of natural cells [20] [21] [22-25], simple water-in-oil emulsions are intrinsically easier to assemble and more stable to environmental change (especially heating) as compartment analogs of cells.

As with standard cells, compartments can maintain a link between the structures of molecules inside the droplet (the “phenotype”) and the structures of the molecules that encode them (the “genotype”). This is important when experimentalists seek to direct the evolution of nucleic acids and their encoded proteins under selective pressure. For example, compartmentalization in two-phase artificial cell systems has been used for directed evolution [3] in the form of “compartmentalized self replication” (CSR). Developed by Tawfik and Griffiths, CSR holds proteins and genes together in water droplets suspended in oil emulsions [26]. CSR was applied to the directed evolution of DNA polymerases by Holliger and his collaborators [10]. CSR has also been used to select DNA polymerases that accept unnatural nucleotides [12] [27].

Compartmentalization is of great interest to synthetic biologists, who seek to put alien “xeno” nucleic acids (xNAs) into artificial cells. One example of xNA are artificially expanded genetic information systems (AEGIS) that increase the number of independently replicable nucleotides from 4 to as many as 12 [28]. By shuffling hydrogen bonding patterns of the G:C and A:T pairs while retaining their Watson-Crick size-complementarity [28]. AEGIS DNA can evolve to produce aptamers molecules that bind to liver and breast cancer cells [29] and isolated proteins [30]. Modified nucleic acids can potentially be used as therapeutics [31]; recently some AEGIS components have been suggested as potential inhibitors of the RNA polymerase from SARS-CoV-2 [32]. The Z:P pair from AEGIS used in this work is shown in Fig 1.

**Figure 1.**
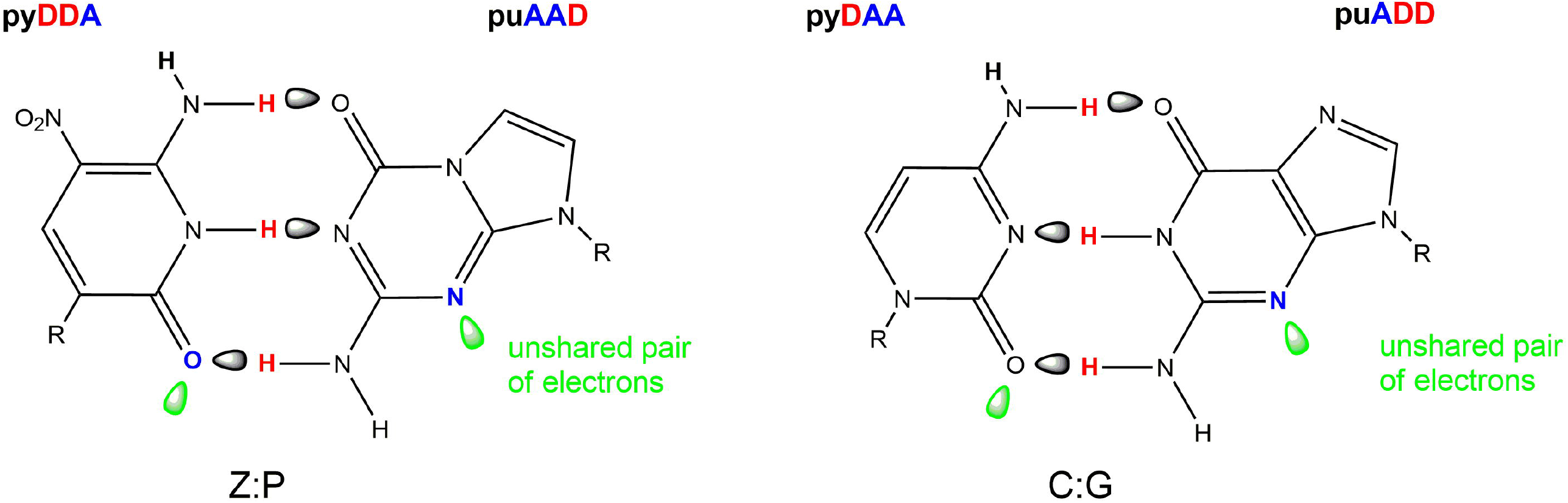
The Z:P pair from an Artificially Expanded Genetic Information System (AEGIS), left. AEGIS nucleotides follow the geometry and hydrogen bonding architecture of the natural nucleotides. The hydrogen bonding patterns are named pu or py, depending on whether they are presented on a large (purine-like) or small (pyrimidine-like) heterocyclic ring system respectively. Hydrogen bond donor (D) and acceptor (A) groups listed starting in the major groove and ending in the minor groove. Unshared pairs of electrons presented to the minor groove are shown in green orbitals. The Z:P pair presents the pyDDA:puAAD hydrogen bonding pattern, while the analog C:G pair presents the pyDAA:puADD. R is deoxyribose.

Compartmentalization in water-in-oil droplets requires a fluid immiscible with water and a surfactant. The surfactant must lower the surface tension of the liquid, stabilize the emulsion, and diminish coalescence [33] [34]. There are several commercially available surfactants and fluids for this purpose; however, there are some difficulties with these. For instance, in our laboratory, Span® 80 has been used to produce water-in-oil droplets, but we have known for years that there are observable differences between batches of the same product (S1 Fig.) potentially arising from the fact that it is derived from animal fat. In our experience, directed evolution experiments using Span80^®^ proved unreliable. We have observed dark precipitates at the bottom of bottles of Span80^®^ and in oil phases with Span80^®^-based mixtures (S2 Fig.). Furthermore, the same authors who pioneered the use of Span® 80 for creating water-in-oil droplets have reported the presence of contamination with peroxides [35] [36], which interferes with some biochemical processes. To counter these issues, silicone fluids were proposed as an alternative. There are many examples of research done with silicone fluids that come from the cosmetic industry. Some, commercialized by Degussa (now Evonik), are not available commercially in research-sized amounts. One cannot expect that a batch of a silicone fluid to be used for producing sunscreen would go through testing for its suitability for PCR. For example Diehl *et al*. report the use of ABIL WE09 from Degussa for water-in-oil emulsions but notes in the supplementary section that dark precipitates are observed [37]. Not surprisingly some researchers fail to replicate the protocols published by other groups or at least need to modify some of the protocols [1]. The lack of more choices of surfactants for biotechnology have motivated other researchers to create their own biocompatible surfactants as well [38].

There are several commercially available fluorinated fluids, which are used for microfluidics. Fluorinated fluids have peculiar properties; they are immiscible with water and hydrocarbons, forming a third “flourous” phase. As natural proteins, membranes, and other biomolecular systems exploit both hydrophilic and hydrophobic phases, a purely hydrophobic oil or surfactant can disrupt biological systems because the hydrophobic portion of the surfactants can invade the hydrophobic cores of proteins and other biologics, disrupting their function. The fluorous phase comprising fluorinated species is sufficiently “orthogonal” to both hydrophilic and hydrophobic systems as to render that third phase inert with respect to most biological molecules systems containing them. Thus, fluorinated oils are used for drug delivery [39, 40], blood substitutes [41, 42], and diagnostics [43]. Several fluorinated oils and surfactants are now commercially available for water-in-oil emulsions, and several of these are widely used. These, however, have densities considerably higher than water, so the water droplets float, which makes it difficult to manipulate them for instance for re-injection. For example, the fluorinated oil HFE-7500 has a density of ∼ 1.6 g/mL [44] [45]. Fluorinert^®^ FC40 has a density between 1.845 and 1.895 [46] [47] [48]. Concentration of the droplets in the top of the fluid also increases the number of droplet-droplet encounters, which encourages droplet fusion.

These considerations provided the motivation to seek fluorinated oils that have a density close to 1 g/mL, the density of aqueous solutions in which biological processes act. The synthesis of silicones from hexamethylcyclotrisiloxane by ring-opening polymerization has been reported before [49]. We report here the synthesis, characterization of polysiloxanes, prepared by ring-opening polymerization under conditions that produce fluids with densities near 1 g/mL while having enough fluorinated monomers to be able to dissolve non-ionic fluorinated surfactants that enable their use for the encapsulation of biochemical reactions.

## Materials and methods

### Materials

Hexamethylcyclotrisiloxane, 1,3,5-trimethyl 1,3,5-tris(3,3,3 trifluoropropyl) cyclotrisiloxane, and 2,4,6,8-tetramethylcyclotetrasiloxane were purchased from Changzhou Fluoride Chemical Co. Genuine Technology. Hexamethyldisiloxane was purchased from Acros. Amberlyst® 15 hydrogen form and potassium trichloro(ethylene) platinate (II) hydrate were purchased from Sigma-Aldrich.

Nonafluorohexyldimethylchlorosilane, 3,3,3 trifluoropropyldimethylchlorosilane; allyloxy(polyethylene oxide) (4-7 EO); allyloxy(polyethylene oxide), methyl ether (6-8 EO) were obtained from Gelest. All chemicals were used without purification. Oligos containing artificial nucleotides, artificial triphosphates dZTP, dPTP were obtained from Firebird Molecular Biosciences. Phi29 polymerase and Exonuclease I were purchased from New England Biolabs. CircLigase™ was purchased from Lucigen. Snap-Cap Microcentrifuge Biopur™ Safe-Lock™ or Safe-Lock Tubes 2 mL were from Eppendorf™. Stainless steel balls, 6 mm were obtained from IKA. EvaGreen® Dye 20X in water was from Biotium. For DNA origami: a “Folding kit basic p7249 Cuboid with large aperture” was obtained from Tilibit Nanosystems. All natural DNA oligos and fluorescent probes were obtained from IDT.

### Ring-opening polymerization

Polysiloxanes were prepared by the ring-opening reaction shown on Fig 2. Two different protocols were developed:

**Figure 2.**
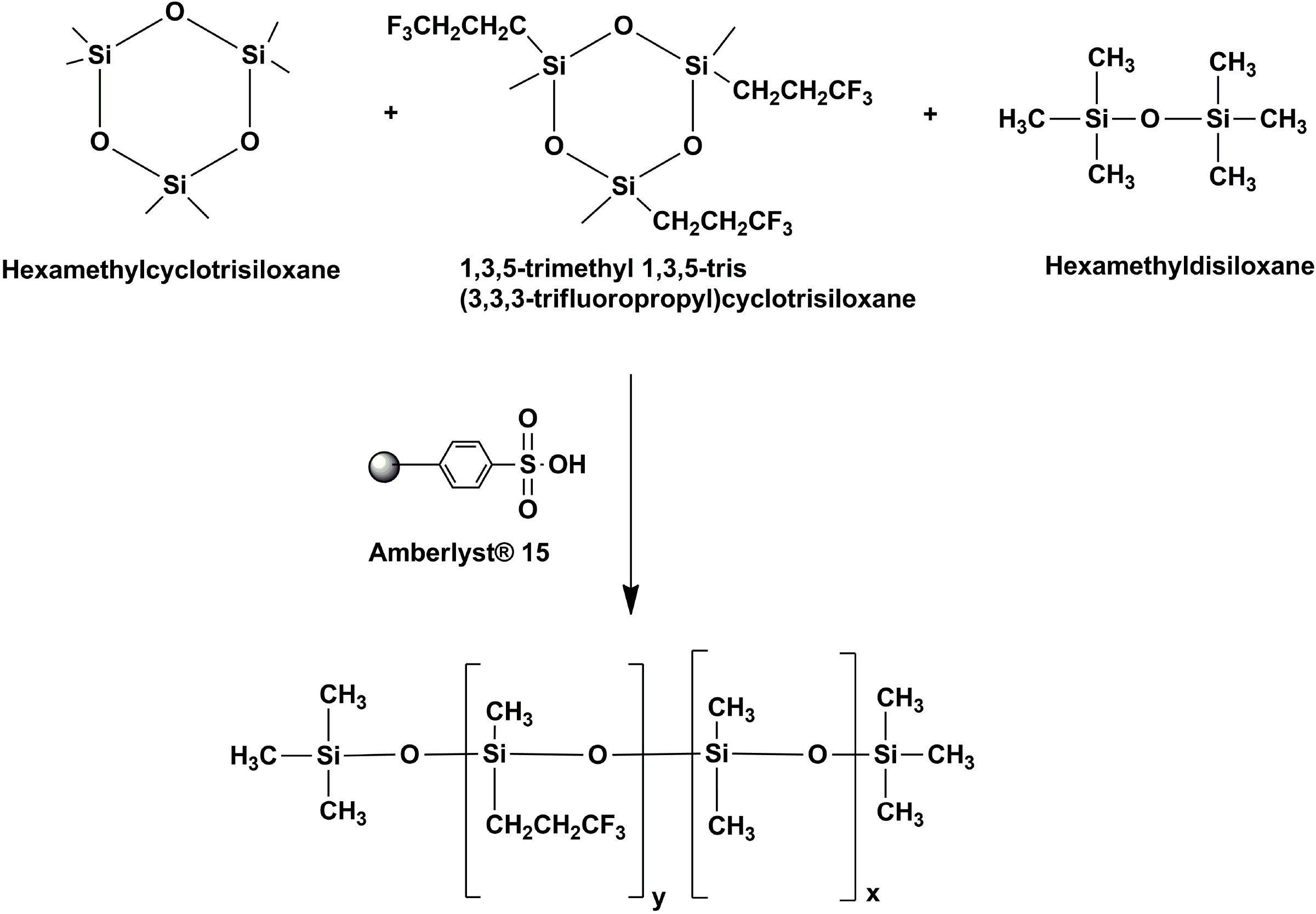
Ring-opening polymerization used to produce the fluorosilicone fluid. The two cyclotrisiloxane monomers (hexamethylcyclotrisiloxane and 1,3,5-trimenthyl 1,3,5-tris(3,3,3-trifluoropropyl)cyclotrisiloxane) are reacted along with hexamethyl disiloxane, which acts as a chain terminator. The catalyst Amberlyst® 15 acts as an acid catalyst for the ring-opening reaction.

#### Protocol 1

The fluorinated monomer 1,3,5-trimethyl 1,3,5-tris-(3,3,3-trifluoropropyl)-cyclotrisiloxane (12 g; 25.6 mmol) was mixed with the non-fluorinated monomer hexamethylcyclotrisiloxane (72 g; 323.7 mmol) and hexamethyl disiloxane (3 mL; 14.1 mmol); the second acted as a chain terminator. The mixture was heated to 75 °C in an oil bath. Amberlyst^®^ 15 catalyst (1.2 g) was added once the reaction mixture was homogeneous. The reaction proceeded under argon for 3 hours at 75 °C. Once cooled, the reaction mixture was decanted from the solid Amberlyst^®^ catalyst.

The products were then resolved by distillation under reduced pressure (23 mm Hg). Unreacted materials were removed at 90 ºC while collecting a fraction of the fluorinated fluid. After removing materials at 90 °C; the remaining fluid was labeled as Fluid A. From this fluid a low molecular weight (LMW, Fluid B) fraction was obtained by distillation (150 °C, 23 mm Hg); a high molecular weight fraction remained, non-distilled (HMW fraction).

#### Protocol 2

The fluorinated monomer 1,3,5 trimethyl 1,3,5-tris(3,3,3-trifluoropropyl) cyclotrisiloxane (25.26 g; 53.9 mmol) was mixed with the non-fluorinated monomer, hexamethylcyclotrisiloxane (36 g; 161.8 mmol) and hexamethyl disiloxane (2.8 mL; 13.11 mmol). The mixture was heated to 75 °C in an oil bath. Catalyst Amberlyst^®^ 15 (1.2 g) was added last, again after the mixture was homogeneous. The reaction proceeded under argon for 4 hours at 75 °C. Then, an opaque liquid was decanted to separate it from the solid catalyst. Distillation using a Kugelrohr apparatus (90 °C, 23 mm Hg) removed unreacted starting materials. This remaining fluid was left at room temperature for 8 hours, after which it had separated into two layers each with different characteristics (top layer: fluid C; bottom layer: fluid D).

### Fluorinated surfactants

Fluorinated hydrophilic chains were synthesized following the work of Kobayashi and Owen [50]. Two fluorinated hydrophobes have been explored: 3,3,3-trifluoropropyl (Pr_*f*_) and 3,3,4,4,5,5,6,6,6-nonafluorohexyl (Hx_*f*_). These hydrophobes were combined with allyloxy(polyethylene oxide) following reference [50] with some modifications. The surfactants (Fig 3) were prepared by hydrolysis of their chlorosilane precursors and eventually bound to the hydrophile by hydrosilation with allyloxy(polyethylene oxide). A scheme of the synthesis and details are provided in the supplementary information section (S3 Fig.).

**Figure 3.**
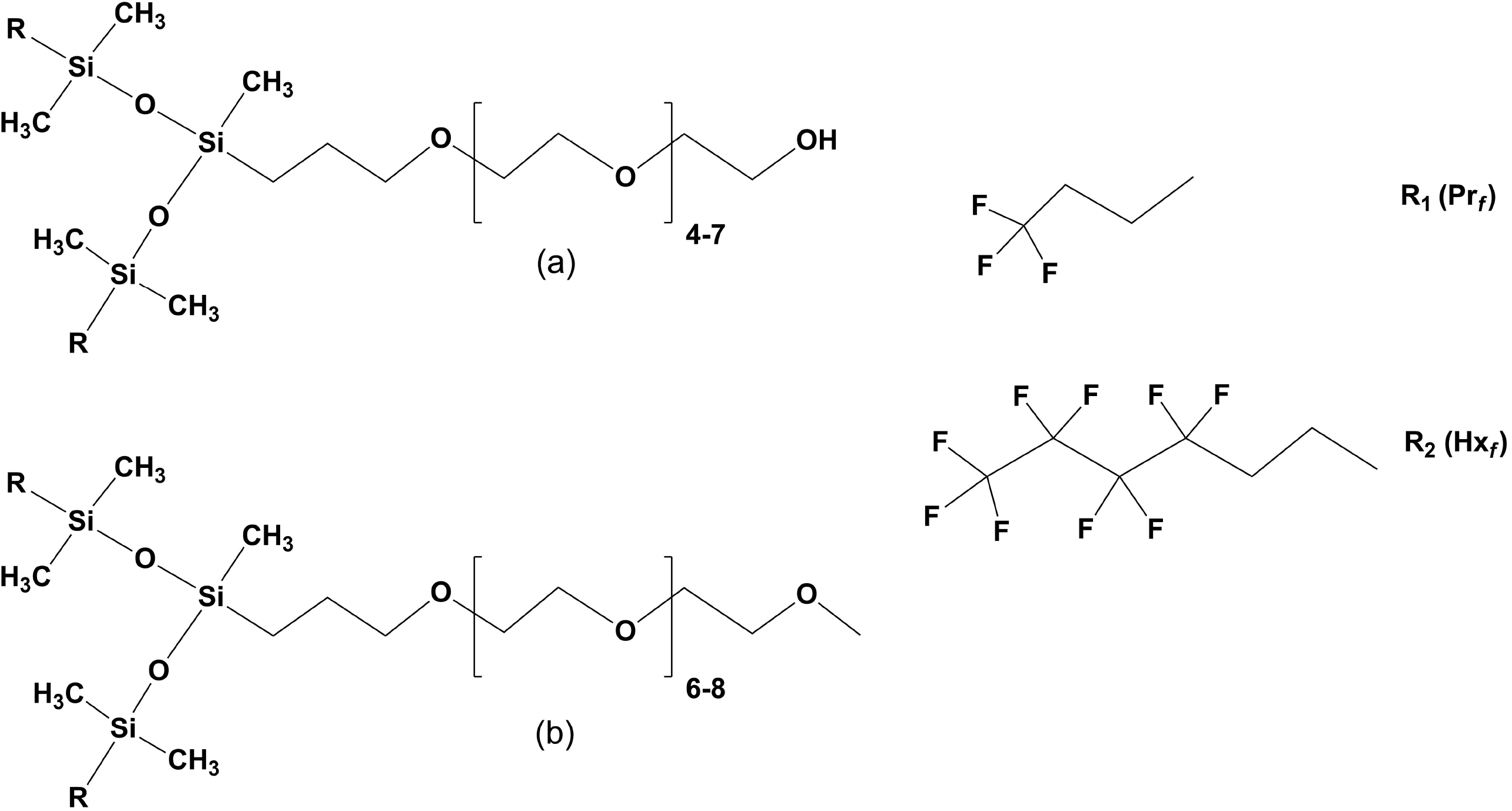
Nonionic fluorosilicone surfactants. The amphiphilic molecules are produced by reacting SiH functionalized branched disiloxanes hydrophobes with allyloxy(polyethylene oxide) following reference [50]. Two different hydrophiles (a, b) and two hydrophobes are shown: one with the 3,3,3-trifluoropropyl (R_1,_ Pr_*f*_) and 3,3,4,4,5,5,6,6,6-nonafluorohexyl (R_2,_ Hx_*f*_).

### Gel permeation chromatography

Gel permeation chromatography (GPC) was performed by Cambridge Polymer Group (Boston, MA) using an Agilent 1100 Series HPLC Gel Permeation Chromatogram equipped with an autosampler, thermostatted column oven, and refractive index detector. The column used for the separation was the Agilent PLgel Mixed-B column (10 μm particle size; 300 mm X 7.5 mm) which has a nominal linear molecular weight separation range from 500-10,000,000 g/mol. These columns are packed with a stationary phase consisting of ∼10 μm gel particles which are composed of a highly crosslinked polystyrene/divinylbenzene matrix. These columns are organic GPC columns, compatible with most organic mobile phase solvents.

Data collected by refractive index (RI) detection was used for characterization of the molecular weight distribution, molecular weight moments, and polydispersity of the samples. The data were analyzed using the Agilent GPC/SEC software v A.02.01. A calibration curve was prepared using polystyrene standards with molecular weights between approximately 500 and 10,000,000 g/mol. A measured amount of the material was transferred to a glass vial with a PTFE lined screw cap. An amount of toluene sufficient to produce a sample concentration of 5 mg/ml was added to each sample, which were then mixed using gentle agitation at a temperature of 30 °C overnight. All samples were clear and colorless. The samples were filtered through 0.45 μm PTFE syringe filters into autosampler vials, filled to the shoulder. Three autosampler vials were prepared of the samples for triplicate injection purposes. Molecular weight moments and polydispersity index (PDI) were calculated by Agilent GPC/SEC Software (v A.02.01) as:

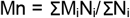

Where Mn is the number molecular weight; M_i_ is the molecular weight of a chain and N_i_ is the number of chains of that molecular weight.

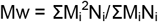

Mw is the weight average molecular weight. In this molecular weight value the more massive chains contribute more to the molecular weight average.

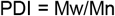

PDI is the polydispersity index.

### Origami DNA

The origami folding mixture comprised 208 staples, each 200 nM (Table in S1 Table) and a scaffold (20 nM), which was a single-stranded DNA p7249 isolated from M13mp18 (7,249 nucleotides) in Tris buffer (5 mM, pH ∼ 8.2, with 1 mM EDTA, 20 mM MgCl_2_, and 5 mM NaCl.

Droplet formation was done using PDMS radial chips that empty droplets of aqueous phase from 20 μm channels into an annular oil flow channel of 50 μm, as previously described by Paegel and Joyce [10]. The continuous phase was a 2% (v/v) of a fluorinated surfactant (a-R_2_) in Fluid B. The continuous phase was pumped at a rate of 5 μL/min while the aqueous phase was pumped at a rate of 7 μL/min.

The origami mixture, both bulk (as a positive control) and emulsified, were placed in a thermal cycler at 65 °C for 10 minutes and then cooled to 60 °C at 0.1° C/s. After this hold, the temperature was lowered by 0.5 °C steps (0.1 °C/s rate) and held at the new temperature for 30 minutes until 40 °C was reached. At the end of the cooling steps the emulsion was centrifugated for 5 minutes at 5,000 xg. Then, 25 μL of the origami reaction were mixed with the provided 6X loading buffer provided with the kit and loaded in an agarose gel 0.5 X Tris-borate EDTA buffer with ethidium bromide and 6 mM MgCl_2_.

### Six-nucleotide PCR with Artificially Expanded Genetic Information Systems (AEGIS)

An aqueous solution holding all components required for a PCR was prepared, consisting of: (a) primers (20 nM each) 5’- GACGGACTGCCTATGAG-3 and 5’- GAGGCGATCGCAGTATC-3’; (b) double stranded template (2 × 10^™7^ μM): 5’- GACGGACTGCCTATGAGCAGTT**Z**AAAAG**Z**TATGATACTGCGATCGC CTC-3’ and 5’- GAGGCGATCGCAGTATCATA**P**CTTTT**P**AACTGCTCATAGGCAGTCCGTC-3’; (c) substrates, dNTPs, dZTP, and dPTP (0.2 mM each), (d) Evagreen 2X; (e) a variant of *Taq* DNA polymerase fused to the DNA binding protein of *Sulfolobus solfactaricus* Sso7d-*Taq*(Δ1-280) [R587Q E626K I707L E708K A743H E832G]) at 7.1 × 10^™2^ μM; and (f) buffer, with 20 mM Tris-HCl, pH 8.4; 10 mM (NH_4_)_2_SO_4_; 10 mM KCl; 2 mM MgSO_4_; 0.1% Triton® X-100.

The PCR mixture (100 μL) was emulsified along with 300 μL of a fluorinated fluid with density 1.009 g/mL that contained 2% (v/v) surfactant (a-R_2_). The fluid was prepared by mixing 1 mL of fluid B with 4 mL of the HMW fluid (protocol 1). The mixture of two phases was placed in a Safe-Lock Tube with and a 6 mm stainless steel bead. The tube was then fitted in a 96-well storage plate and secured between the top and bottom plates of TissueLyser II (QIAGEN). The components were then mixed for 10 seconds at 15 Hz followed by 7 seconds at 17 Hz. Then, 50 μL of emulsion were transferred to PCR tubes with an optical lid by triplicate. The PCR used the following cycle: 92 °C for 30 seconds 40 cycles: [92 °C for 10 seconds; 48 °C for 15 seconds; 72 °C for 20 seconds] and acquired fluorescence after each cycle using a Lightcycler^®^ 96 (Roche Life Science).

### Taqman ® assay in droplets

An aqueous solution containing the components of a Taqman^®^ assay reaction was prepared to contain:

- Template: 5’-CAGCAGTGCCAGCAGAACAAAGGTATTCATCTTAGTGA**P**GTGCGC AGTCAGCTCACTACT-3’, at 0.4 μM concentration.
- Primer: 5’-AGTAGTG AGCTGACTGCGC-3’, at 0.2 μM concentration; and
- Probe: 5’-FAM-ATGAATACC-ZEN-TTTGTTCTCTGCTGG CAACTGCTG-3’

The other components of the reaction mixture were: (a) substrate dNTPs and dZTP, each at 0.2 mM; (b) buffer at 10 mM Tris pH 8.3, 1.5 mM MgCl_2_, 50 mM KCl; and (c) enzyme, a variant of *Taq* DNA polymerase [R587Q, E832C], 6 pmol/50 μL reaction.

The aqueous mixture (100 μL) was emulsified along with 300 μL of a fluorinated fluid with density 1.009 g/mL that contained 2%(v/v) surfactant (a-R_2_). The fluid was prepared by mixing 1 mL of fluid B with 4 mL of the HMW fluid (protocol 1). The tube containing the two phases was then fitted in a 96-well storage plate and secured between the top and bottom plates of TissueLyser II (QIAGEN). The components were then mixed for 10 seconds at 15 Hz followed by 7 seconds at 17 Hz. Then, 50 μL of emulsion were transferred to PCR tubes with an optically clear lid and placed in the following cycle: 40 cycles of: [95 °C for 8 seconds; 62 °C for 24 seconds] and acquired fluorescence after each cycle using a Lightcycler^®^ 96 (Roche Life Science).

### Rolling circle amplification (RCA) in droplets

An aqueous mixture comprising all of the components for a RCA reaction was prepared to include: 1X Phi29 buffer (New England Biolabs), which has 50 mM Tris-HCl at pH 7.5, 10 mM MgCl_2_, 10 mM (NH_4_)_2_SO_4_, and 4 mM DTT (@ 25 °C); (b) triphosphates, dATP, dCTP, dTTP, dGTP, 0.2 mM each; primer at 0.8 μM, having the sequence 5’-CAGGGCTGGGCATAGAAGTCAGGGCAGA-3’; polymerase, Phi29 0.1 U/μL, with BSA 0.2 μg/μL; circular ssDNA template molecule at 2.5 μg/μL. The circular ssDNA template was prepared by circularizing: P-5’TATGCCCAGCCCTGTA AGATGAAGATAGCGCACAATGGTCGGATTCTCAACTCGTATTCTCAACTCGTAT TCTCAACTCGTCTCTGCCCTGACTTC-3’ with CircLigase™ (Lucigen) according to the manufacturer protocol. Non-circularized material was degraded with Exonuclease I (*E. coli*) according to the manufacturer protocol as well.

The aqueous mixture holding all the components necessary for RCA was emulsified by placing aqueous mixture (100 μL) along with 300 μL of a fluorinated fluid with density 1.009 g/mL that contained 2%(v/v) surfactant (a-R_2_). The fluid was prepared by mixing 1 mL of fluid B with 4 mL of the HMW fluid (protocol 1). The tube containing the two phases was then fitted in a 96-well storage plate and secured between the top and bottom plates of TissueLyser II (QIAGEN). The components were then mixed for 10 seconds at 15 Hz followed by 7 seconds at 17 Hz. Then, 50 μL of emulsion were transferred by triplicate to PCR tubes with an optically clear lid. Fluorescence was acquired after each minute using a Lightcycler^®^ 96 (Roche Life Science).

### Expression and purification of the DNA polymerase variants

A plasmid (pJExpress414, DNA 2.0) containing the gene encoding the *Taq* DNA polymerase variant of interest was expressed in *E. coli* C43(DE3) (37 ºC, overnight growth, IPTG 0.5 mM) in LB media (50 mL). The next morning, cells were pelleted (∼ 0.75 g) in a refrigerated centrifuge for 10 minutes at 5,000 x g and stored at -20º C. The pellet was resuspended in 2.5 mL of Tris 50 mM; glucose 50 mM; EDTA 1 mM; pH 8.0 and heat-shocked at 75 ºC for 10 minutes, then mixed with 3.75 mL of BugBuster® master mix (Novagen) for 20 minutes at room temperature. The lysate was pelleted in 1.5 mL tubes, at 21,000 xg for 5 minutes and the supernatant was mixed with NiNTA resin (1.5 mL of resuspended resin washed and resuspended in 4 mL of Lysis buffer: Tris HCl, 20 mM pH 8.0; NaCl 500 mM, imidazole 10 mM). The lysate and the resin were then incubated and gently spun at 4 ºC for 1 h.

The lysate was then loaded on a column and washed with: (a) 2 mL wash: Tris, 20 mM; NaCl, 1 M; followed by (b) 1 mL wash: Tris, 20 mM; NaCl, 150 mM; followed by (c) 2 mL wash: Tris, 20 mM; NaCl, 500 mM; imidazole, 20 mM; and (d) eluted with elution buffer: 1 mL Tris HCl, 10 mM, pH 8.0, NaCl 250 mM, imidazole 500 mM. Buffer was exchanged by ultracentrifugation (AMICON, 15 mL, 50,000 MW cutoff) to final storage buffer, which was 10 mM Tris pH 8.0; 100 mM KCl; 0.5 mM DTT; 0.1 mM EDTA; 0.5% Tween 20; 0.5 % Igepal; glycerol 50%. Protein concentration was measured using Pierce™ BCA Protein Assay Kit, following the manufacturer recommendations.

## Results

Fluorine has an atomic mass of 19, meaning fluorinated molecules are intrinsically denser than water. However, this greater atomic mass can be partly offset by the larger mean volumes of CF_2_ (38 Å^3^) and CF_3_ (92 Å^3^) groups relative to CH_2_ (27 Å^3^) and CH_3_ (54 Å^3^) [31], both less dense than water. Therefore, we reasoned that fluorous oils having a density of water might be obtained by balancing their fluorinated components with alkane components. This balancing must also consider the density of silicon (Si = 28 g/mole) since cyclotrisiloxanes are part of the polymerization mixture.

With these considerations, we explored methods to prepare low-density fluorous oils by reacting siloxanes having both alkane and trifluoromethylalkane side chains, specifically 1,3,5-trimethyl 1,3,5-tris(3,3,3-trifluoropropyl)-cyclotrisiloxane and hexamethylcyclotrisiloxane (Fig 2). It was taken into consideration that the growth of the polymeric chain must be limited since high molecular weight molecules will increase the viscosity of the fluid. For this purpose the chain termination component (hexamethyldisiloxane) plays an essential role.

### Ring-opening polymerization

Acid-catalyzed ring-opening polymerization following protocol 1, produced polysiloxanes with bimodal molecular weight distribution (Fig 4). Protocol 2 produced two fluids with different densities which spontaneously separated. Interestingly, each of these fluids exhibits a bimodal molecular weight distribution as well (Fig 5). Following protocol 1, fluid A was separated by distillation using a Kugelrohr apparatus. Obtaining 15 mL of the lower molecular weight fraction (150 ºC, 23 mm Hg) with a density of 1.0565 g/mL. The remaining fluid (∼ 53 mL) consists of a high molecular weight fraction with density 1.031 g/mL; this remained undistilled up to 200 ºC at 23 mm Hg). Table 1 lists molecular weight values for different fractions.

**Table 1.** Summary of molecular weight moments and polydispersity.

| sample | Mp [g/mol] | Mn [g/mol] | Mw [g/mol] | PDI |
| --- | --- | --- | --- | --- |
| Fluid A | 9,404 ± 57.7 | 1748 ± 31.2 | 10,126 ± 83.5 | 5.793 ± 0.058 |
| Fluid A peak 1 | 10,321 ± 34.6 | 3,206 ± 486 | 11,478 ± 74.0 | 3.637 ± 0.561 |
| Fluid A peak 2 | 321 ± 2.08 | 230 ± 2.65 | 305 ± 2.00 | 1.326 ± 0.008 |
| Fluid C | 6,412 ± 64 | 2,064 ± 11 | 6,266 ± 116 | 3.04 ± 0.06 |
| Fluid C peak 1 | 6,412 ± 64 | 4,549 ± 34 | 7,529 ± 144 | 1.66 ± 0.019 |
| Fluid C peak 2 | 626 ± 6 | 597 ± 3 | 637 ± 3 | 1.07 ± 0.008 |
| Fluid D | 645 ± 4 | 1,722 ± 12 | 6,543 ± 119 | 3.80 ± 0.07 |
| Fluid D peak 1 | 7,178 ± 47 | 4,892 ± 23 | 8,522 ± 70 | 1.74 ± 0.009 |
| Fluid D peak 2 | 645 ± 4 | 611 ± 4 | 661 ± 4 | 1.08 ± 0.003 |

**Figure 4.**
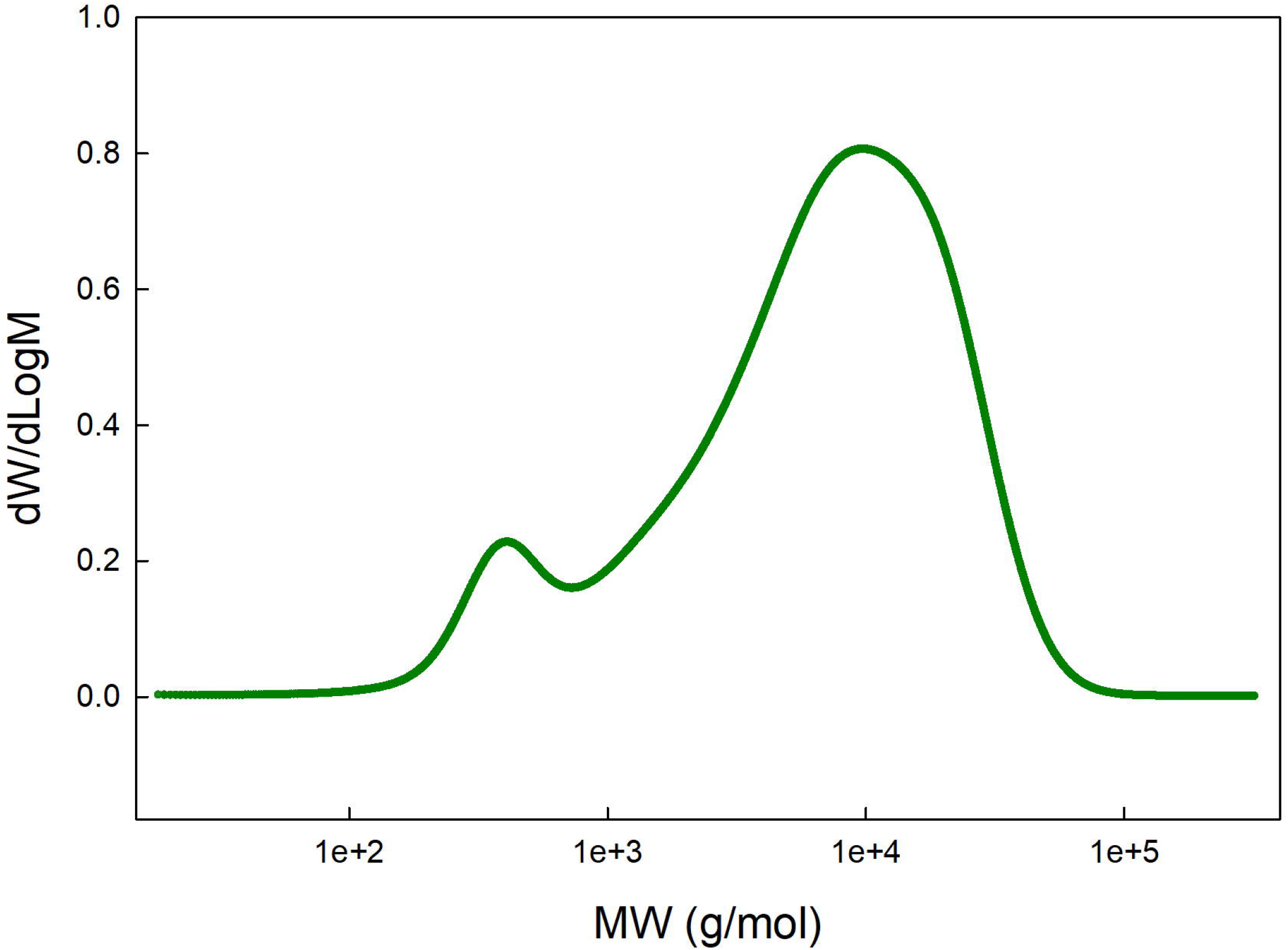
Molecular weight distribution of polysiloxanes produced via protocol 1. Ring-opening polymerization following “protocol 1” produced polysilioxanes with bimodal distribution. These two peaks can be separated by distillation under reduced pressure with a high molecular weight peak (Mp ∼ 9,400 g/mol) and a low molecular weight peak (Mp ∼ 300 g/mol). The low molecular weight fraction contains a higher concentration of fluorinated monomers estimated by ^1^H-NMR and is capable of dissolving the fluorinated surfactants. Peak 1 corresponds to the higher molecular weight peak (rightmost) and peak 2 corresponds to the lower molecular weight peak (leftmost). S4 Fig. shows an overlay of two gel permeation chromatography runs of the two distinctive fractions separated by distillation.

**Figure 5.**
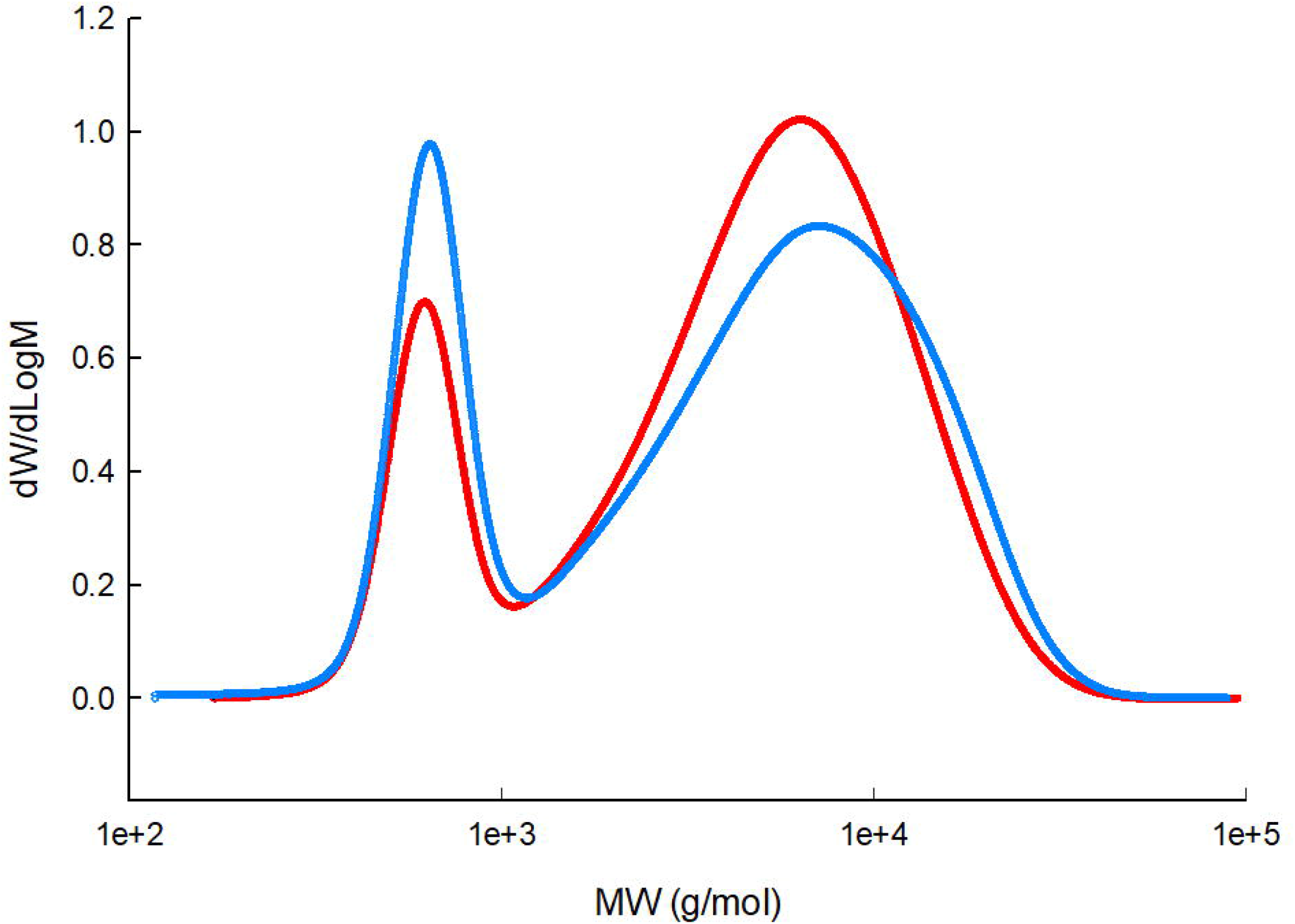
Molecular weight distribution of polysiloxanes produced via protocol 2. Ring-opening polymerization following “protocol 2” produced two liquid polysilioxanes that spontaneously separate in two phases with different densities and are referred as fluid C (red) and fluid D (blue). Each fluid exhibits bimodal molecular weight distribution.

Reported molecular weight moments and polydispersity from the overall chromatogram, as average +/- one standard deviation. Molecular weights are reported in polystyrene equivalents. Fluid A peak 1 is referred as high molecular weight fraction (HMW) and Fluid A peak 2 referred as low molecular weight fraction (LMW). Mn = ΣM_i_N_i_/ΣN_i_, where Mn is the number molecular weight; M_i_ is the molecular weight of a chain and N_i_ is the number of chains of that molecular weight. Mw = ΣM_i_^2^N_i_/ΣM_i_N_i,_ where Mw is the weight average molecular weight. In this molecular weight value the more massive chains contribute more to the molecular weight average. PDI is the polydispersity index. PDI = Mw/Mn.

#### ^1^HNMR analysis

Resonances with chemical shifts between 0.0 and 0.3 ppm were assigned to Si-CH_3_ (labeled “a” in Fig 6). These signals arise from both fluorinated and non-fluorinated monomers. Resonances with chemical shifts between 0.7 and 0.9 ppm were assigned to the methylene groups attached to Si (α-CH_2_-, labeled “b” in Fig 6). Resonances with chemical shifts between 1.9 and 2.2 ppm were assigned to the methylene beta to the Si (β-CH_2_-, labeled ‘“c” in Fig 6) for the fluorinated monomers that contained -CH_2_^α^-CH_2_ ^β^-CF_3_.

**Figure 6.**
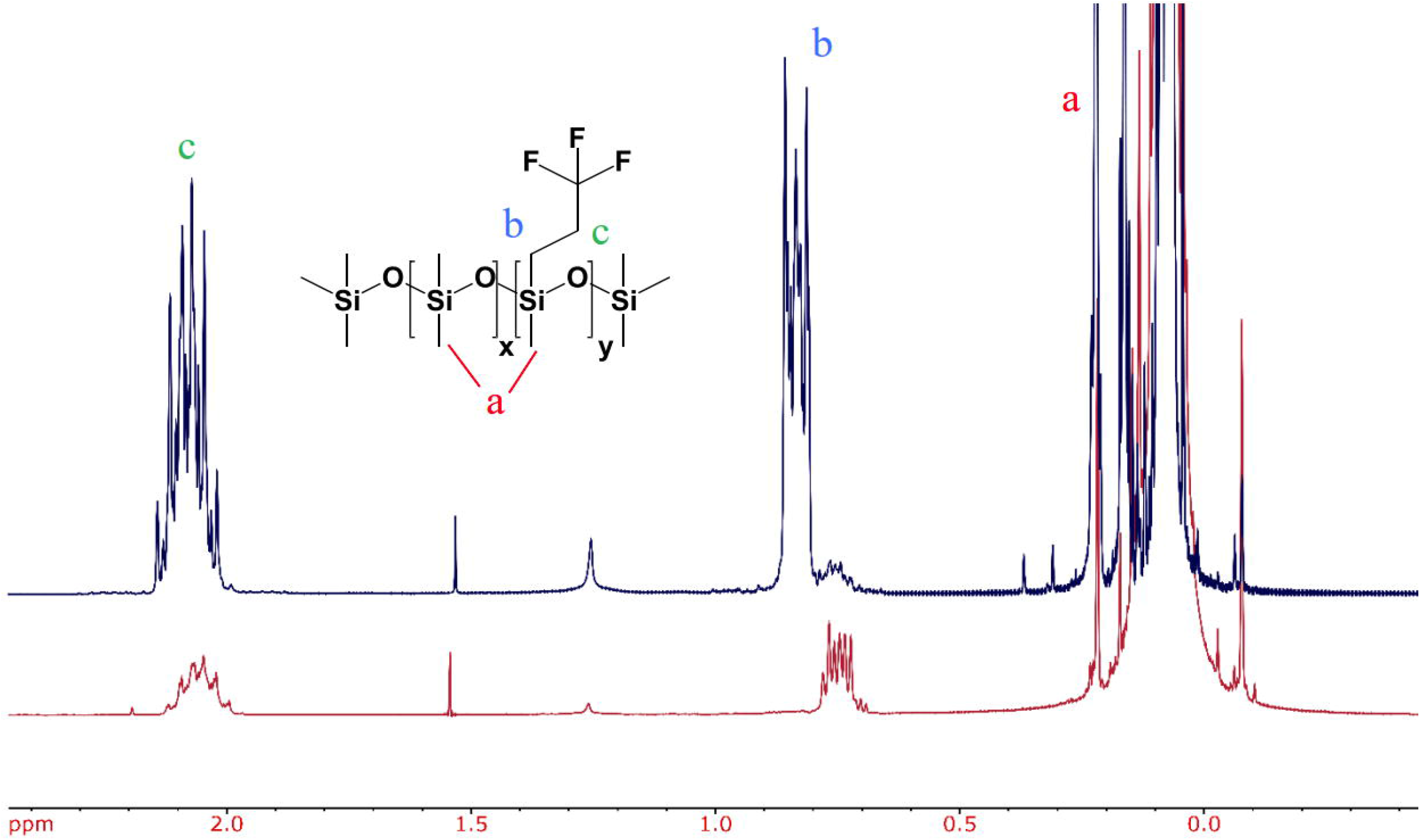
^1^NMR analysis of fluorosilicone polysiloxanes. Overlay of the ^1^HNMR of the low molecular weight (blue) and the high molecular weight (red) fraction samples indicating the assigned peaks.

The ratios of non-fluorinated monomer to fluorinated monomer were calculated using the integrals from the β protons (denoted by “c”) and the Si-CH_3_ protons (denoted by “a”). The following equation was used to determine the monomer ratio:

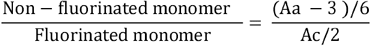

Here, A_c_ is the integration area of the peak for the β-CH_2_-protons and A_a_ is the integration area of the peak for the Si-CH_3_ protons. The integration values and monomer ratios are summarized in Table 2.

**Table 2.**
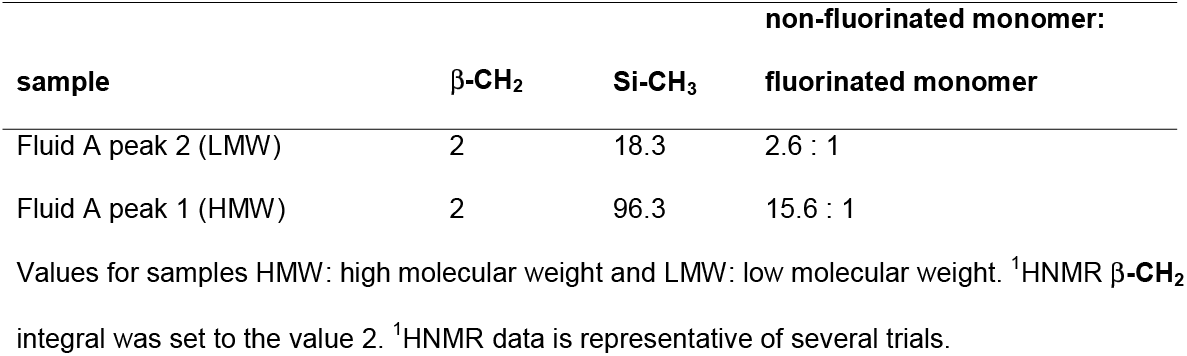
Integration of peaks used for determination of the ratio of non-fluorinated monomer: fluorinated monomer.

The integral of the resonances assigned to the Si-CH_3_ units (0.0 to 0.3 ppm) is inversely proportional to the fraction of fluorinated monomers present in the fluid. It was observed that fractions with higher fluorinated monomers exhibited lower viscosity. As expected, the fractions with a higher degree of fluorination were capable of solubilizing the nonionic fluorinated surfactants. Figs 7 and 8 show ^19^FNMR and ^1^HNMR of homogeneous solutions of the surfactants in the low molecular weight fraction (Fluid B, protocol 1). These solutions remained homogeneous for at least 10 days. The NMR spectra show the distinctive signals that arise from the surfactants solubilized to 4% (v/v). Table 3 shows different values of density, viscosity and proportion of non-fluorinated to fluorinated monomer compositions in the fluorinated polysiloxanes.

**Table 3.** Viscosity, density and other properties measured on fluorinated polysiloxanes.

| sample | density<br>(g/mL) | Viscosity<br>(cSt) | <sup>1</sup>HNMR<br>β-CH<sub>2</sub><br>integral | <sup>1</sup>HNMR<br>Si-CH<sub>3</sub><br>integral | MW<br>(g/mol) | PDI | non-fluorinate<br>fluorinated<br>monomer |
| --- | --- | --- | --- | --- | --- | --- | --- |
| Fluid A peak 2 | 1.031 | n/d | 2 | 18.3 | 305 | 1.3 | 2.55 : 1 |
| Fluid A peak 1 | 1.0565 | n/d | 2 | 96.3 | 11,478 | 3.6 | 15.55 : 1 |
| HMW fluid | 0.986 | 67 | 2 | 58.0 | 10,126 | 5.8 | 9.17 : 1 |
| Fluid C | 1.0457 | 6.06 | 2 | 33.98 | 6,266 | 3.04 | 5.16 : 1 |
| Fluid D | 1.1347 | 10.8 | 2 | 12.24 | 6,543 | 3.8 | 1.54 : 1 |
| Fluid B * | 1.1397 | 4.5 | 2 | 11.48 | n/d | n/d | 1.41 : 1 |

Summarized viscosity and molecular weight analysis for fluorinated polysiloxane samples. Reported viscosity as fitted using a Newtonian model. Determined by Cambridge Polymer Group using a TA Instruments DHR-2 rheometer. Reported molecular weight moments and polydispersity from the overall chromatogram, as average +/- one standard deviation. Molecular weights are reported in polystyrene equivalents. Fluid B\* was prepared following protocol 1 but with a shorter time of reaction obtaining a fluid with lower viscosity but with higher density and higher proportion of fluorinated monomers. <sup>1</sup>HNMR β-CH<sub>2</sub> integral was set to the value 2. Some tests not performed are indicated as “not determined” n/d.

**Figure 7.**
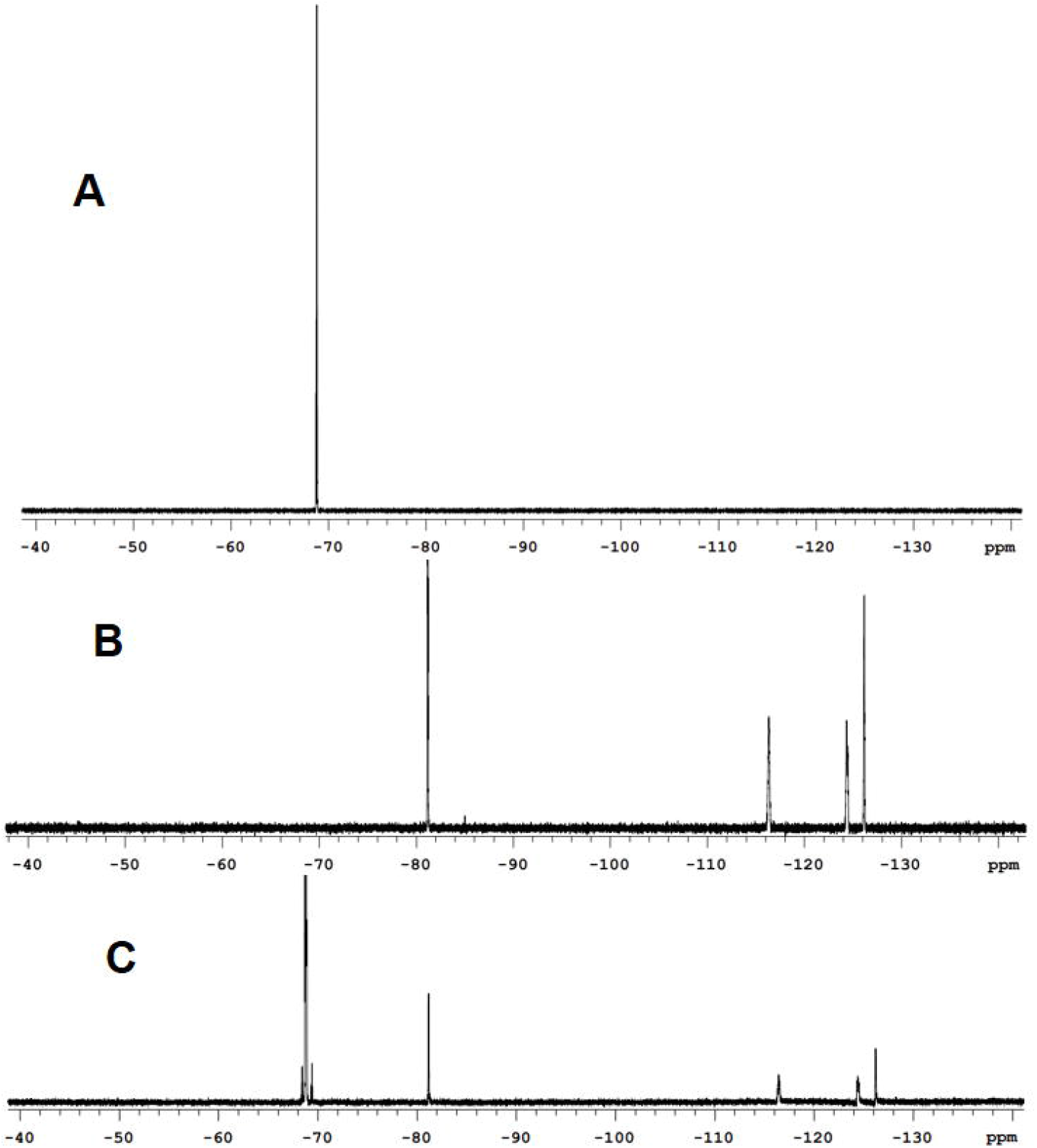
^19^FNMR overlays showing the presence of the surfactant with the Hx_*f*_ hydrophobe CF_3_CF_2_CF_2_CF_2_CH_2_CH_2_- dissolved in the low molecular weight fluid (fluid B). (A) A single peak that comes from the C**F**_3_CH_2_CH_2_-from the fluorinated fluid. (B) Four peaks that correspond to the fluorine of the Hx_*f*_ hydrophobe C**F**_3_C**F**_2_C**F**_2_C**F**_2_CH_2_CH_2_-. (C) ^19^FNMR of a homogeneous 4% solution (v/v) of the surfactant in the fluorinated fluid exhibiting the signals from both components.

**Figure 8.**
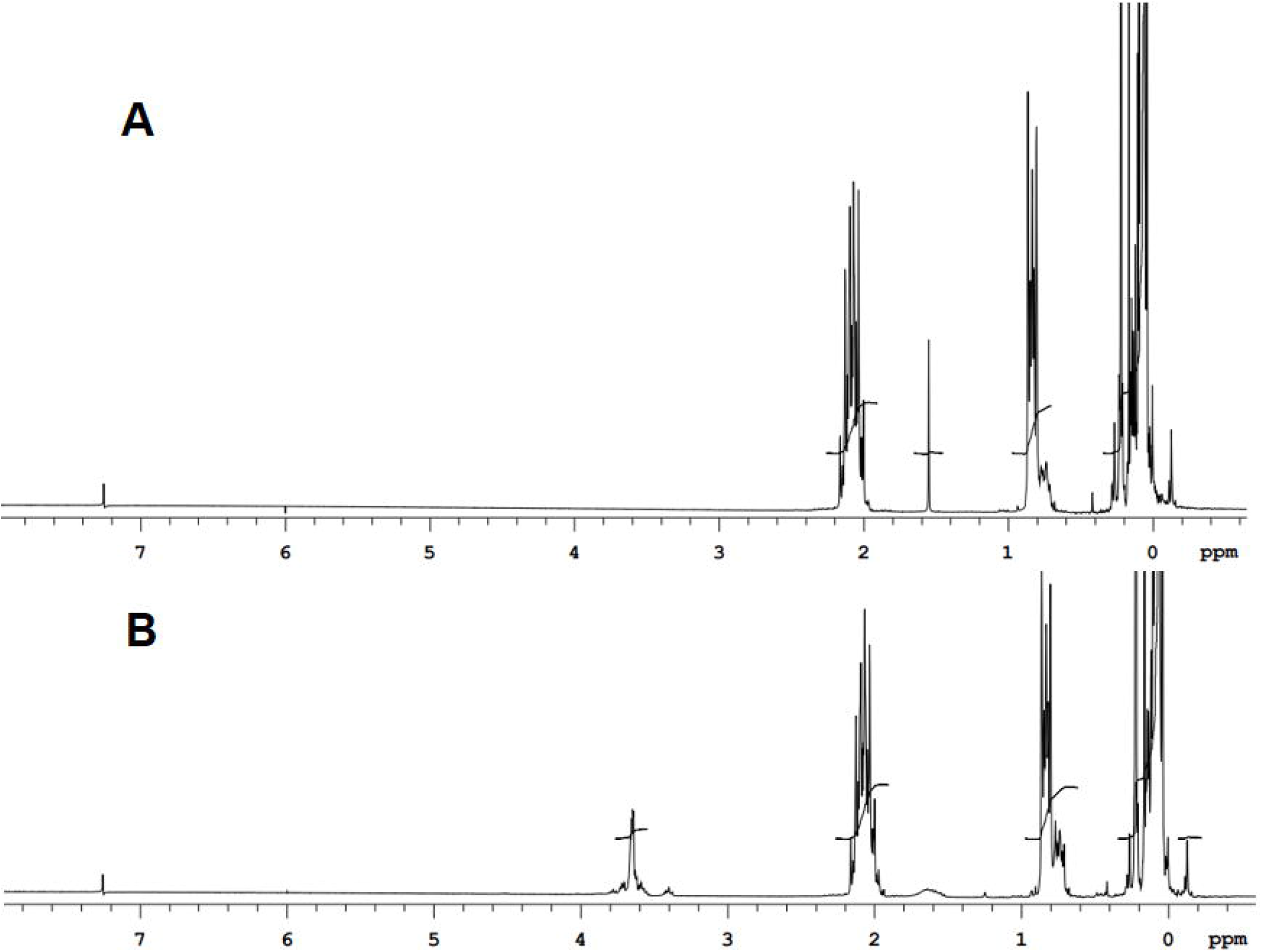
^1^HNMR overlays showing the presence of the surfactant with the Pr_*f*_ hydrophobe CF_3_CH_2_CH_2_- dissolved in the low molecular weight fluid (fluid B). (A) ^1^HNMR of the fluid without surfactant. (B) ^1^HNMR of a 4% (v/v the fluid with surfactant.A single peak that comes from the C**F**_3_CH_2_CH_2_-from the fluorinated fluid. The characteristic signals of the -(CH_2_CH_2_-O)- can be observed at 3.4-3.8 ppm.

### Droplet formation

Droplets shown in this work were made using two methods: using microfluidic devices or by shaking using a QIAGEN TissueLyser II. Fig 9 shows droplets produced with a radial microfluidics chip. Fig 10 shows droplets formed by shaking 300 μL of the fluid/surfactant with 100 μL of the aqueous phase in a Safe Lock tube with a 6 mm stainless steel bead.

**Figure 9.**
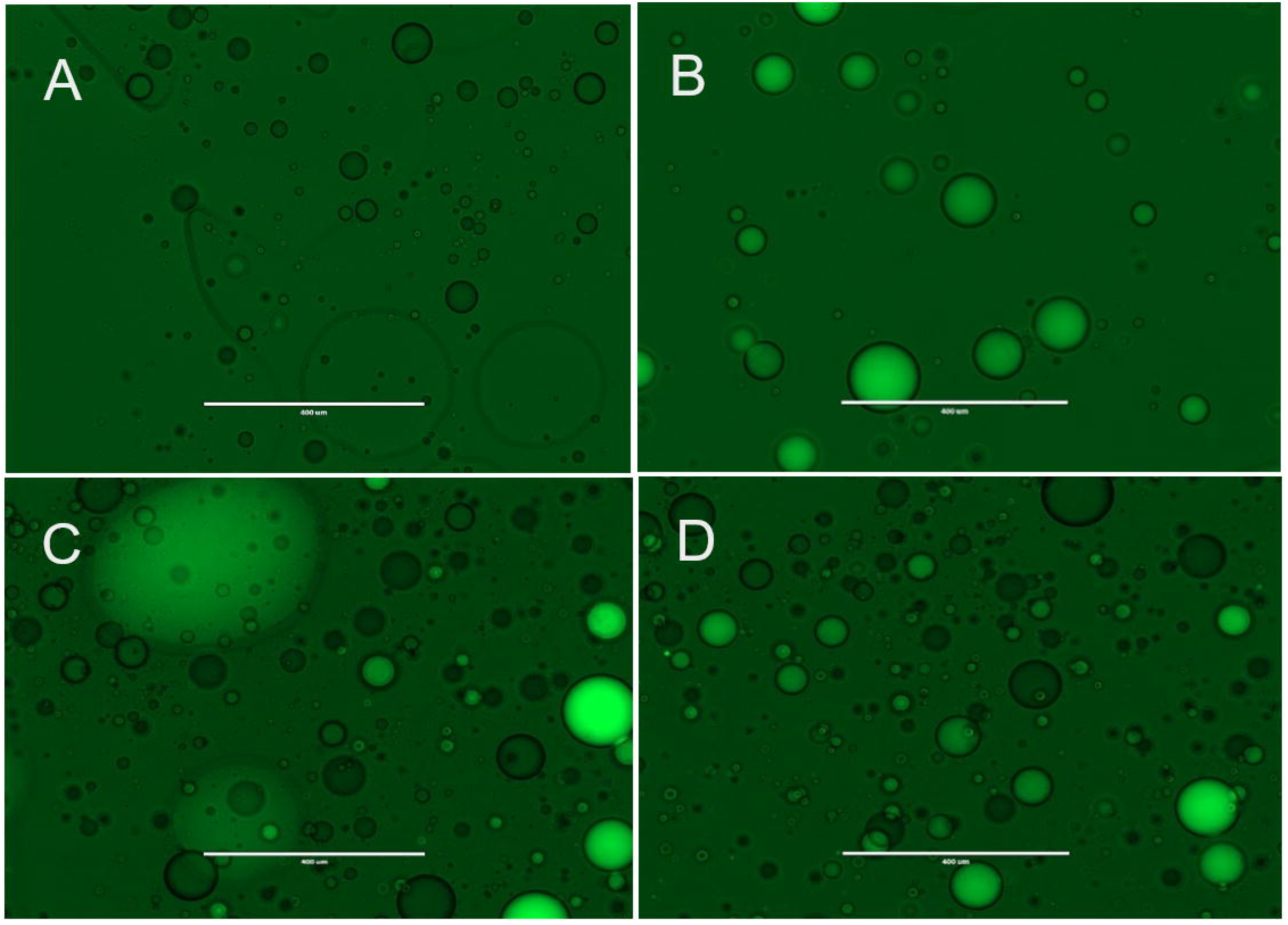
Transmission microscope images of droplets containing oligos with a FAM fluorescent molecule or a Black hole quencher (BHQ). The fluorinated fluid (fluid B) with 1% of surfactant (b-R_1_) was pumped at 10 µL/min. Aqueous phase contained 5 µM of 5’-CTGCTCATAGG CAGTCCGTCA-BHQ-3’ or 5’-FAM-TGACGGACTGCCTATGAGCAG-3’ in 20 mM Tris-HCl, pH 8.4; 10 mM (NH_4_)_2_SO_4_; 10 mM KCl; 2 mM MgSO_4_; 0.1% Triton® X-100. Aqueous phase was pumped at 2 µL/min. (A) Droplets carrying oligo with Black Hole Quencher (BHQ). (B) Droplets carrying oligo with FAM. (C). Droplets carrying BHQ placed on the left side of the glass slide and droplets carrying FAM placed on the right hand side of the slide. (D) After being in contact for 10 minutes no fusion is observed. Scale bar is 400 µm. Pictures are representative of several trials.

**Figure 10.**
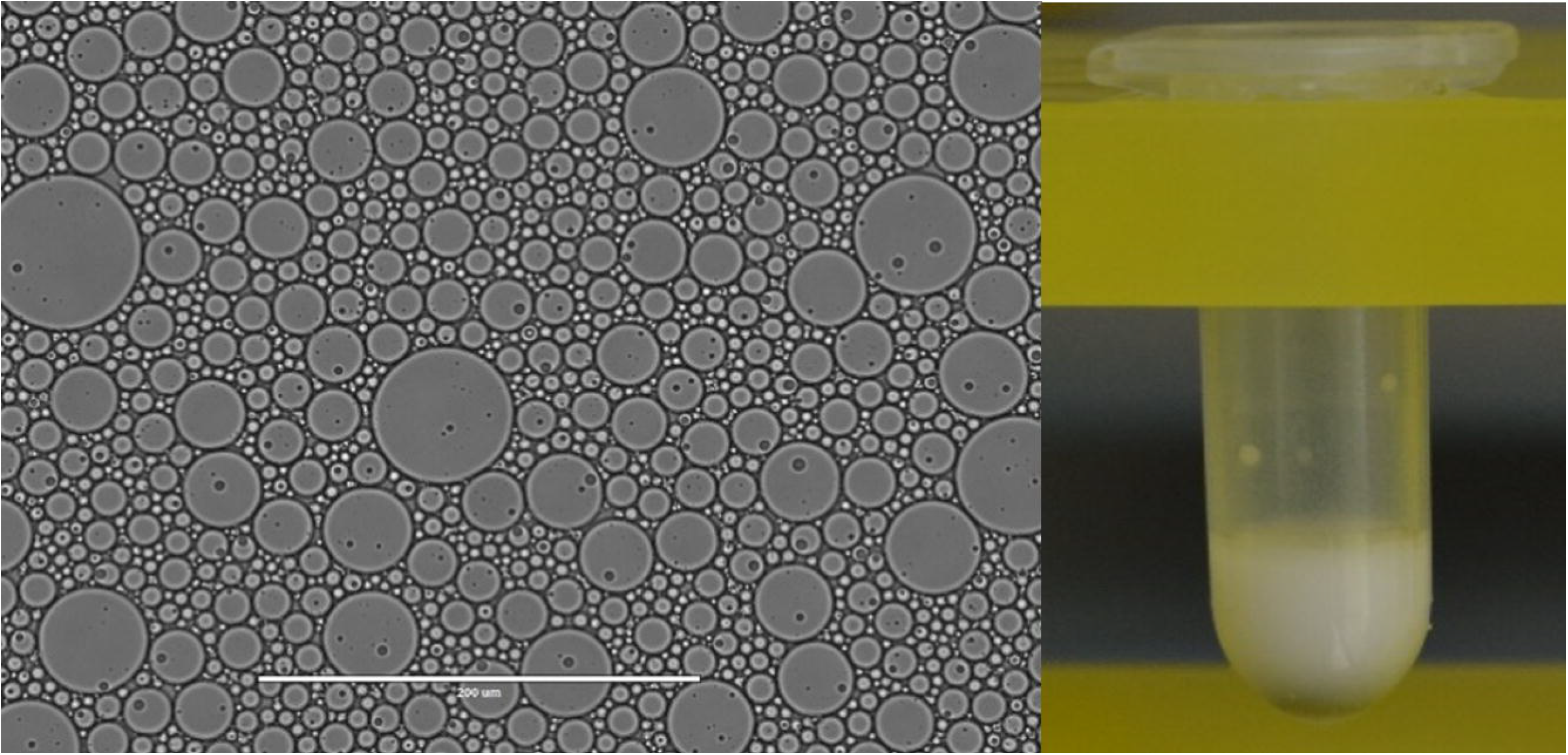
Emulsion made with fluid C (protocol 2). Left: transmission microscope image of droplets. Scale bar is 200 µm. Right: picture of the emulsion formed by shaking 100 µL of buffer and 300 µL of fluid C with 2% of the surfactant a-R_2_ containing the Hx_*f*_ hydrophobe. The emulsion is formed by shaking the contents in a Snap-Cap Microcentrifuge Biopur™ Safe-Lock™ Tube containing a 6 mm stainless steel bead. The tube was fitted in a TissueLyser II and shaken for 10 seconds at 15 Hz followed by 7 seconds at 17 Hz. Scale bar is 400 µm. Pictures are representative of several trials.

Next, we challenged the emulsions prepared with these fluid/surfactant mixtures to host processes that involve both DNA and enzymes, all useful in biomolecular sciences, nanotechnology: (a) Assembly of origami DNA nanostructures inside droplets suspended in the fluorinated oil. (b) PCR amplification of 6-letter DNA templates inside droplets suspended in the fluorinated oil, including those built from an AEGIS DNA alphabet. (c) Generation of fluorescent signals inside droplets suspended in the fluorinated oil using a Taqman® assay, also with six-letter DNA real-time PCR. (d) Generation of fluorescent signals inside droplets suspended in the fluorinated oil from the formation of RCA products.

### Origami DNA

An origami kit obtained from Tilibit Nanosystems that assembles to form a cuboid of dimensions 61 nm x 8 nm x 52 nm with an aperture of 9 nm x 15 nm was set up as suggested by the manufacturer. The formation of the nanostructure was verified by its distinctive shift in migration compared with the scaffold when electrophoresed in a 2% agarose gel in 0.5 X TBE with ethidium bromide and 6 mM MgCl_2_. The migration pattern of the scaffold DNA and a correctly assembled nanostructure is shown in S8 Fig. The purified nanostructure has been characterized by Transmission Electron Microscopy (S9 Fig.). Our results showed that both the emulsified and non-emulsified origami reaction produced a nanostructure with the expected migration pattern (Fig 11). The result shown in Fig 11 comes from droplets prepared using microfluidics as described in the experimental section. This experiment was repeated preparing the emulsions using a TissueLyser II (QIAGEN) without noticeable difference between the methods of emulsification (S5 Fig.).

**Figure 11.**
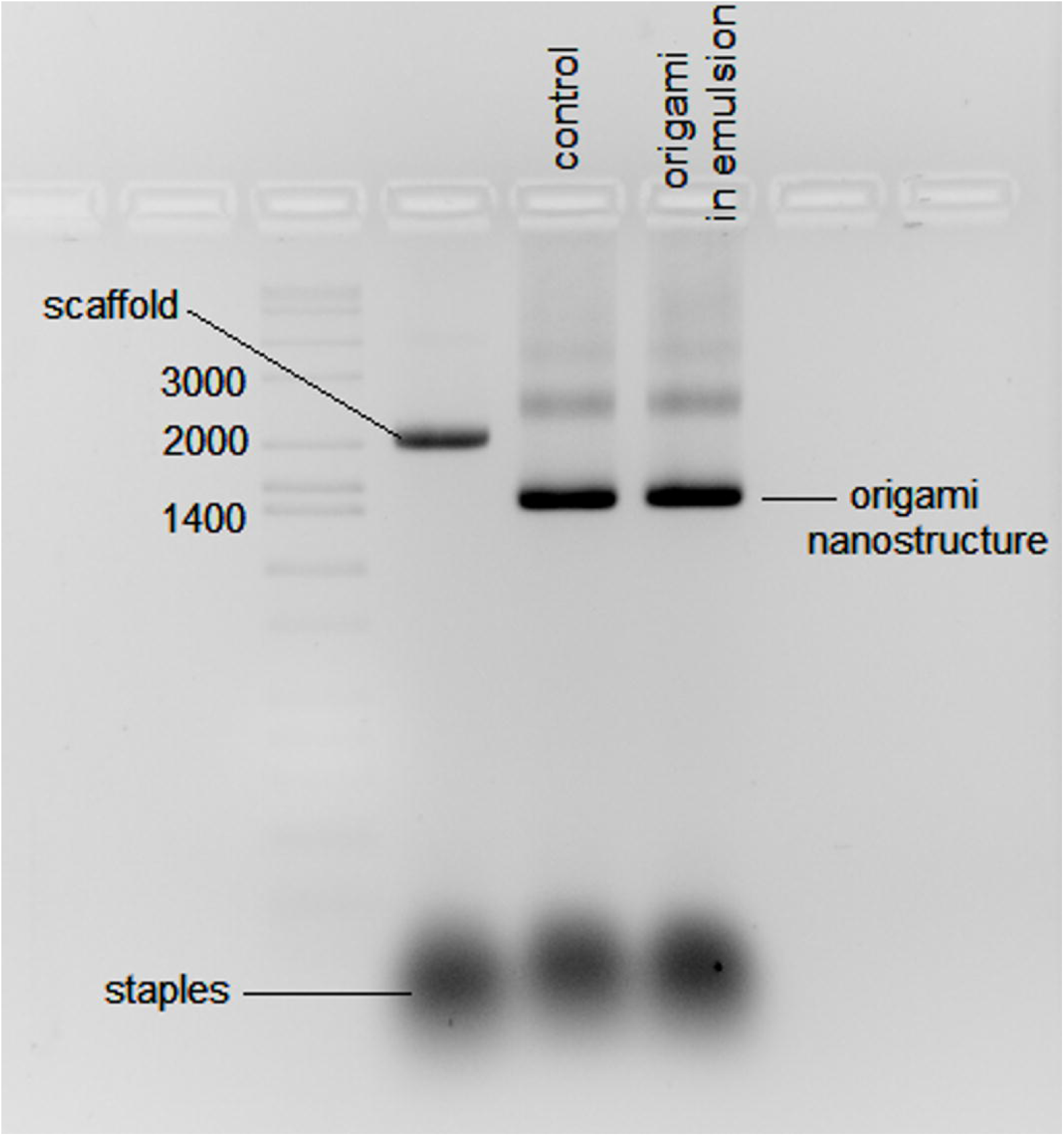
DNA electrophoresis in agarose (2%) showing the formation of origami structures in droplets. The origami products were run in 0.5 X Tris-borate EDTA buffer with ethidium bromide and 6 mM MgCl_2_. The origami structure folded is called “nanostructure PF-2, cuboid with large aperture” from Tilibit Nanosystems. The scaffold, a single-stranded DNA type p7249 isolated from M13mp18 of length 7,249 bases. The staples are a mixture of 208 oligos (S1 Table 1). A sample of the mixture of scaffold and staples not annealed is run for comparison. The origami folded properly shows a distinctive migration on the gel (see Figs S8 and S9 for reference). Annealing was done by incubating at 65 °C for 10 minutes then cooled down from 60 °C to 40 °C at a rate of 1 °C per hour.

### AEGIS PCR with 6-Letters

To demonstrate that droplets suspended in the iso-dense fluorinated oil had thermal stability adequate to support PCR and, in particular, PCR containing AEGIS bases, a double stranded DNA molecule with 49 bp containing the AEGIS pair Z:P (Fig 1) was used as template. The enzyme used to amplify this was a derivative of the *Taq* DNA polymerase where the amino acids 1-280 were deleted (removing the exonuclease domain), with the N-terminus of the catalytic domain fused to a processivity enhancing domain from *Sulfolobus solfataricus*. The *Taq* variant has 6 mutations, two specific for enhancing the incorporation of the pair **Z:P** (R587Q and E832G). The enzyme is described as: Sso7d-*Taq* (Δ 1-280) R587Q E626K I707L E708K A743H E832G.

Progress of the PCR was monitored by observing the fluorescence of Evagreen^®^, which intercalates into double-stranded PCR amplicons. Fig 12 shows real time PCR fluorescence signals arising from the PCR reaction. As a control a PCR done in bulk aqueous solution shows comparable fluorescence signals to the ones that come from water droplets emulsified in iso-dense fluorinated oil. These results showed that neither the surfactant nor the fluorinated oil had any measurable impact on the progress of the PCR. This is consistent with the orthogonality of the fluorinated oil and the surfactant with respect all biological molecules involved in the PCR.

**Figure 12.**
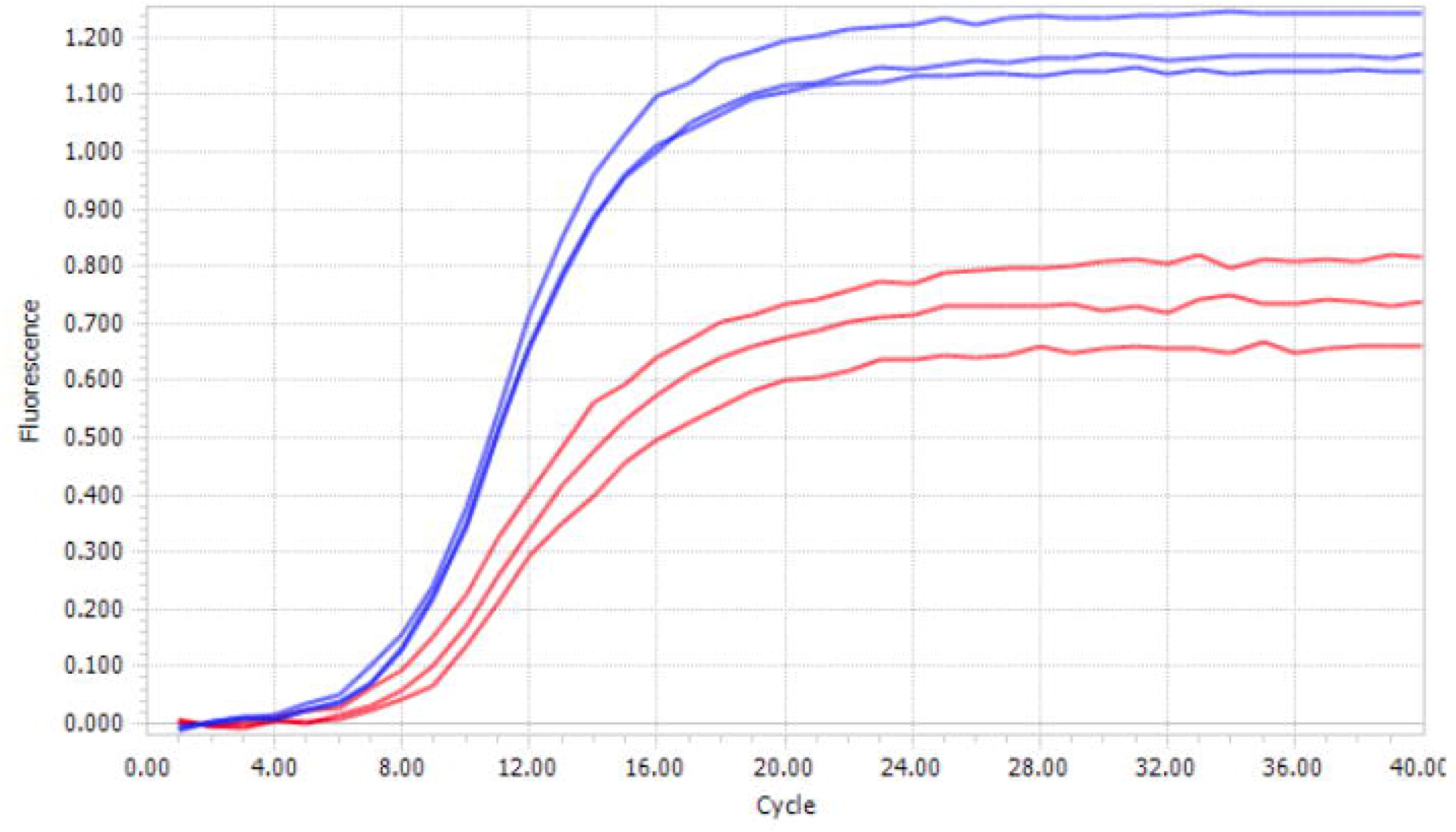
Real time PCR showing emergence of fluorescence from intercalated Evagreen^®^ without emulsification (blue lines) and in the emulsified PCR (red lines). The aqueous phase was emulsified by mixing 100 μL of aqueous phase mixed with 300 μL of fluorinated oil (1 volume of LMW fraction and 4 volumes of the HMW fraction, see protocol 1) with 2% of surfactant (a-R_2_). Assays done by triplicate.

### Taqman^®^ assay

One standard way to detect DNA is a Taqman^®^ assay. Here, DNA polymerase from *Thermus aquaticus* degrades the downstream strand using its 5’-3’ exonuclease activity displacing a probe from the template, cleaving the probe in the process. This cleavage separates a fluorescent reporter (here, fluorescein, FAM) from one end of the probe from a fluorescence quencher at the other end of the probe, to give a fluorescent signal.

Here, we adapted the Taqman^®^ assay to detect a target analyte that contained AEGIS dZ and dP (Fig 1). As before, the aqueous phase was emulsified by mixing 100 μL of aqueous phase mixed with 300 μL of fluorinated oil (1 volume of LMW fraction and 4 volumes of the HMW fraction, see protocol 1) with 2% of surfactant (a-R_2_). Emulsions were made by shaking using a QIAGEN TissueLyser II (15 Hz for 10 seconds followed by 17 Hz for 7 seconds) in a Snap-Cap Microcentrifuge Biopur™ Safe-Lock™ tube containing a 6 mm stainless steel bead. Again, output was essentially the same as observed in bulk Taqman^®^ assays (Fig 13). The *Taq* DNA polymerase variant used contains two mutations (R587Q, E832C) that favor the incorporation of the Z:P pair

**Figure 13.**
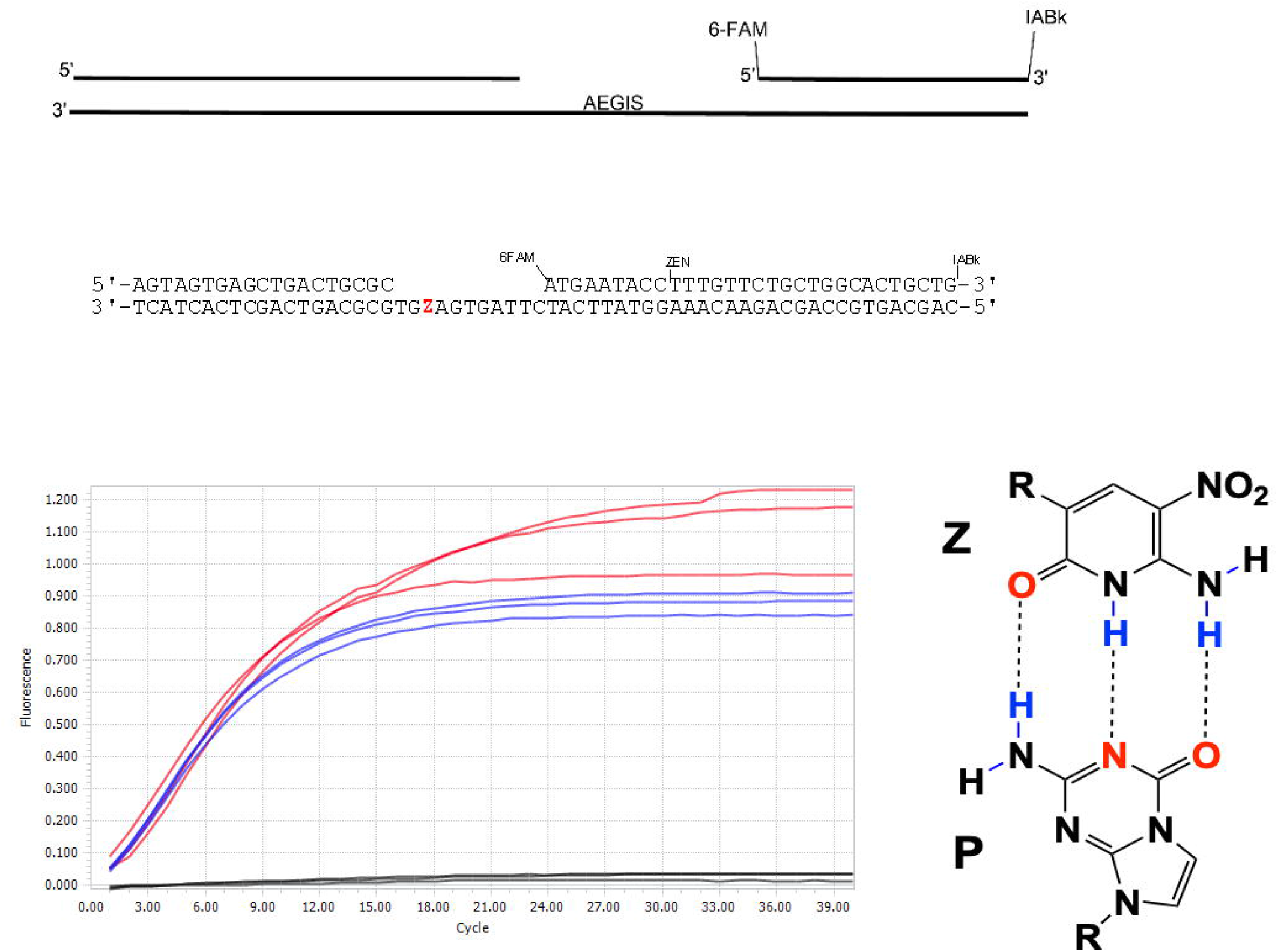
Taqman^®^ assay in droplets. Top: Scheme of the Taqman^®^ assay containing AEGIS (dZ:dP pair hydrogen bonding shown). Bottom: Real-time fluorescence detected from a probe. The probe contains a 5′ fluorophore: 6-carboxyfluorescein (6FAM) and is double-quenched with: 3′ Iowa Black™ (IABk) and a proprietary internal ZEN quencher (IDT). When *Taq* [R587Q E832C] polymerase places a dZ opposite dP and the 5’-3’ exonuclease of *Taq* polymerase cleaves the probe eventually releasing a fluorescent molecule. The blue line is the fluorescence from a non-emulsified reaction. The red line corresponds to the signal from the emulsified reaction. The emulsion was made by mixing 100 μL of aqueous phase with 300 μL of a fluorinated oil with 2% (v/v) fluorinated surfactant as shown on Fig 2. Mixed in QIAGEN TissueLyser II in the presence of a 6 mm stainless steel bead at 15 Hz for 10 seconds followed by 17 Hz for 7 seconds using an Eppendorf Safe-Lock Tube (2 mL) containing a stainless steel ball. Assays were done by triplicate.

### RCA in droplets

RCA works by having a circular template, a primer that binds to that circular template, and a strand displacing polymerase that extends that primer, displacing already synthesized DNA as it proceeds around the circle. Here, the aqueous phase used to prepare the droplets contains a circularized template 96 nucleotides around, a primer complementary to the circular template, and dNTPs. The polymerase was Phi29 DNA polymerase. The progress of the RCA was followed by measuring fluorescence emitted by an intercalating fluorescent dye (EvaGreen®) in a Light Cycler over a period of 1 hour (Fig 14).

**Figure 14.**
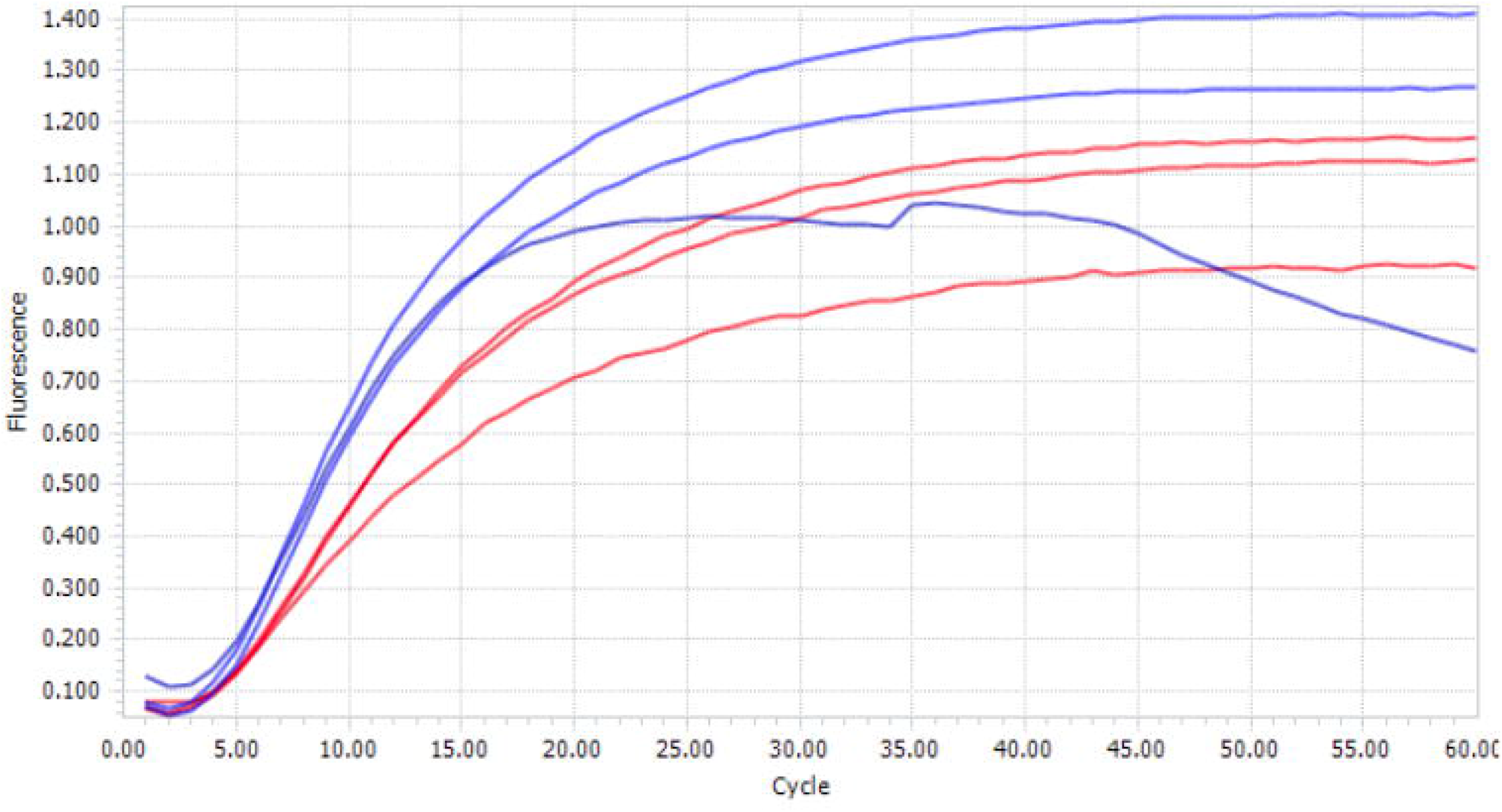
Amplification curves obtained using the LightCycler96 for RCA amplification in droplets. Fluorescence detection (EvaGreen®) is acquired after each two step cycle [37 °C for 30 seconds -37 °C for 30 seconds followed by aquisition] x 60 cycles. Blue lines correspond to the reaction that was not emulsified. Red lines correspond to the emulsified reaction. Assays done by triplicate.

## Discussion

The growing interest in synthetic cells [51] matches a growing interest in engineering enzymes for synthetic biology using directed evolution [52, 53] [54]. For example, Tawfik and Griffiths [26] developed a method to link genotype and phenotype in droplets of water emulsified in oil called “Compartmentalized Self-Replication” (CSR). This strategy was adapted in the Holliger laboratory for the directed evolution of DNA polymerases [10].

However, various laboratories have reported difficulties in reproducing CSR procedures. One of the first problems identified was associated to the surfactant Span 80^®^ [55]. It is very likely that Span 80^®^, which is derived from animal fat, carries in uncontrolled amounts a problematic contaminant. Indeed, the Holliger group reports the need for an antioxidant to avoid oxidation due to peroxide contaminants in Span 80^®^ [35]. Thus, while results are possible with surfactant mixtures containing Span 80^®^, these are hit-or-miss, and has become known in the community as problematically irreproducible.

For this reason, many have turned to synthetic materials and a third liquid phase, the “fluorous phase”. Kobayashi and Owen reported some time ago the synthesis of non-ionic fluorosilicone surfactants with 3,3,3-trifluoropropyl and 3,3,4,4,5,5,6,6,6-nonafluorohexyl side chains with poly(oxyethylene) hydrophiles [50].

Here, balancing the mixture of hydrocarbon-carrying silanes, intrinsically less dense than water, with fluorohydrocarbon-carrying silanes, intrinsically more dense than water, yielded mixtures that had densities essentially identical to the density of water. As the density of water can change with different amounts of solutes, the density of the fluorinated oils described here can be changed by mixing the fractions, separated by distillation, in different ratios to match the desired density.

The fluorinated oils described here have been used to encapsulate different biochemical reactions such as DNA origami folding, PCR, Taqman^®^ assay and RCA. We found no negative impact of the encapsulation. All of these biochemical reactions worked as well in these iso-dense fluorinated oil-surfactant systems as their non-emulsified analogs. Thus, they add a significant and important new resource for nanotechnology, directed evolution and synthetic biology.

## Acknowledgments

We thank Tamara Aigner and Jean-Philippe Sobczak from Tilibit Nanosystems for useful discussions and assistance in troubleshooting and confirming the formation of the origami nanostructure. We thank Brian Paegel laboratory for providing molds and assistance in preparing microfluidic devices as well as valuable discussions in surfactant design.

## Supporting information

**S1 Table 1. Sequences of oligos used as staples for origami structure PF-2 cuboid with large aperture**.

**List of oligos used as staples in Origami DNA folding experiment**.

**S1 Figure. PCR in emulsion using different batches of Span**^**®**^ **80**.

**S2 Figure. Dark precipitates observed in Span**^**®**^ **80**.

**S3 Figure. Example of synthesis of fluorinated nonionic surfactant a-R**_**2**_.

**S4 Figure. Overlay of molecular weight distribution of the two fractions separated by distillation after ring-opening polymerization following protocol 1**.

**S5 Figure. Origami fold in droplets using a QIAGEN TissueLyser II**.

**S6 Figure. Raw image of S5 Figure**.

**S7 Figure. Raw image of Fig 11**.

**S8 Figure. DNA electrophoresis of cuboid with large aperture from Tilibit Nanosystems**.

**S9 Figure. Transmission Electron Microscopy (TEM) of a correctly assembled DNA origami cuboid with large aperture from Tilibit Nanosystems**.

**S10 Figure. Droplets made using a microfluidic device (T-junction)**.

**S1 File. MW distribution. Molecular weight distribution data for Fig 4 and S4 Fig**.

**S2 File. MW distribution. Molecular weight distribution data for Fig 5**.

